# The Kifc3 Motor Protein Controls Centrosomal Factor Cep192 in Ontogenic Coordination of Megakaryocyte Development

**DOI:** 10.64898/2026.03.20.713234

**Authors:** Kamaleldin E. Elagib, Sijie Liu, Valentin Burguener, Ranjit Sahu, Deepika M. Kotay, Ciorsdaidh Watts, Gerhard Wolber, Adam N. Goldfarb

## Abstract

The distinct features of neonatal megakaryocytes, high proliferation and inefficient platelet production, have clinical repercussions. A diminished capacity for stress thrombopoiesis, the response to acute drops in platelet counts, contributes to the high prevalence of thrombocytopenia in premature infants and to impaired platelet recovery after umbilical cord blood stem cell transplantation. High proliferation also promotes leukemogenesis in babies with Down Syndrome (DS). The transcriptional coactivator Mkl1/MrtfA participates in programming the ontogenic shift from fetal/neonatal to adult-type megakaryopoiesis; in this activity it is opposed by the DS-associated kinase Dyrk1a. In a screen for downstream ontogenic effectors in human progenitors, we identified the kinesin Kifc3 as a factor selectively decreased in adult megakaryocytes and whose knockdown in neonatal megakaryocytes induced adult-type morphogenesis with augmented platelet release. Kifc3 acts as a minus-end directed motor for centrosomal delivery of various cargos. Centrosomal release of Cep192 has recently been found induce cellular process extensions through actin remodeling, reminiscent of megakaryocyte platelet release. In our studies, Cep192 showed striking upregulation and dispersion in adult vs neonatal megakaryocytes, and Kifc3 knockdown recapitulated this effect in neonatal megakaryocytes. A role for Cep192 in promoting megakaryocyte morphogenesis, distinct from its role in centrosome biogenesis, was demonstrated *in vitro* and *in vivo*. *In silico* screening for Kifc3 inhibitors identified a small molecule that affected neonatal megakaryocytes similarly to Kifc3 knockdown, indicating feasibility for therapeutic argeting of the Kifc3-Cep192 pathway in clinical conditions associated with fetal-type megakaryopoiesis.

**Key Points:**

1. The motor protein Kifc3 dictates megakaryocyte ontogeny in association with its control of the centrosomal actin-remodeling factor Cep192.
2. Knockdown or small molecule targeting of Kifc3 enhances neonatal megakaryocyte morphogenesis and thrombopoiesis.

## Introduction

As compared with adult-type megakaryocytes (Mk), fetal-type Mk display increased progenitor proliferation, decreased morphogenesis, and decreased platelet release^1^. Mk morphogenesis comprises a coordinated sequence of events that primes the cells for intravascular platelet release: endomitosis, polyploidization, cellular enlargement, and polarized process extension. Impairment in this program occurs most prominently during fetal development but persists in normal infants for up to a year and strongly manifests in iPSC-derived megakaryocytes^1–3^. These ontogenically controlled features ensure functional adaptation to host needs. Thus, overall development of the fetus is best supported by rapid Mk expansion and by production of hypoactive platelets^1^. However, fetal-type megakaryopoiesis lacks the capacity for stress thrombopoiesis, leading to clinical problems in situations requiring rapidly augmented platelet production^1^. Two relevant scenarios consist of neonatal thrombocytopenia, a condition affecting a high proportion of premature infants^4^, and impaired platelet recovery after umbilical cord blood hematopoietic stem cell transplantation^5,6^. The high proliferation rate of fetal-type Mk also contributes to the high rate of neonatal neoplasms in the setting of Down syndrome^7^. Finally, the sparse platelet production by fetal-type Mk constitutes a major bottleneck in the translation of *ex vivo* platelet production to routine clinical application^8,9^. Understanding the mechanisms that control the ontogenic transition from fetal to adult-type megakaryopoiesis will enable rationale design of therapies for these clinical problems.

Both cell intrinsic and environmental factors influence Mk ontogeny^1,10^. Cell intrinsic differences encompass signaling and transcriptional properties, the former including thrombopoietin responsiveness, mTOR activation, and p21 levels^1,11^. Cell intrinsic differences in transcription arise from the fetal-specific expression of the RNA-binding protein Igf2bp3 which prevents induction of the coactivator Mkl1/MrtfA, a master regulator of adult-type Mk morphogenesis^12,13^. The deficiency of Mkl1/MrtfA in fetal-type Mk can be overcome by driving its nuclear translocation through Dyrk kinase inhibition, triggering an ontogenic switch to adult-type morphogenesis and platelet production^2^. However, this approach lacks therapeutic feasibility due to the broad involvement of Dyrk kinases in basic cellular functions and the tumor suppressor activity of Dyrk1a in fetal-type Mk^14–16^. Discovery of ontogenic effectors downstream of Dyrk-Mkl1/MrtfA will thus expand opportunities for clinical intervention while also elucidating the machinery orchestrating adult-type Mk morphogenesis.

Experiments in this study identified the motor protein Kifc3 as an Mk ontogenic mediator downregulated in the fetal-adult transition. Kifc3 decline initiated a novel pathway in which upregulation and dispersion of the centrosomal protein Cep192 enabled remodeling of actin cytoskeleton from fetal-type cortical banding to adult-type cytoplasmic sheets. A critical role for Cep192 in adult Mk morphogenesis was confirmed *in vitro* by knockdown in human progenitors and *in vivo* by CRISPR mutation in mice. Prior design of small molecule inhibitors of the paralog Kifc1^17^ supported the feasibility of Kifc3 targeting. Computational modeling of small molecule docking on the motor domain of Kifc3 yielded a compound that recapitulated the effects of Kifc3 knockdown in fetal-type Mk: enhanced morphogenesis, increased platelet production, Cep192 upregulation and dispersion. These results offer a novel treatment approach to fetal-Mk associated problems and implicate Cep192 in coordinating the orderly execution of adult Mk morphogenesis.

## Methods

### Cell culture

Cryopreserved neonatal cord blood (CB) and adult peripheral blood (PB) CD34+ primary human progenitors were purchased, thawed, pre-stimulated, and subjected to unilineage Mk, erythroid, or granulocytic culture as previously described^12,18^. The Dyrk kinase inhibitors harmine and EHT 1610 were obtained, dissolved in DMSO, and applied to cultures as recently indicated^2^. Centrinone was purchased from Selleckchem (cat # S7837), dissolved in DMSO, and used at a concentration of 125 nM. For *ex vivo* cultures of murine progenitors, fetal liver samples underwent erythrocyte depletion with Gibco^TM^ ACK lysing buffer (ThermoFisher) followed by washing with cold PBS. To expand progenitors, cells were cultured one day in RPMI-1640 with 10% FBS, supplemented with murine cytokines from Peprotech (50 ng/ml SCF, 50 ng/ml Interleukin-6, 20 ng/ml Interleukin-3), as well as 50 μM 2-mercaptoethanol, 100 U/ml penicillin, and 100 U/ml streptomycin. For Mk differentiation cytokine composition was changed to 40 ng/ml human TPO plus 25 ng/ml murine SCF, and cultures continued for an additional 3 days.

### Cell transduction and transfection

Lentiviral constructs for shRNA-mediated knockdowns of *KIFC3* and *CEP192* consist of the pLKO.1 vectors produced by The RNAi Consortium (TRC): clone IDs TRCN0000116462 (*KIFC3* 62), TRCN0000116463 (*KIFC3* 63), TRCN0000135907 (*CEP192* 1), TRCN0000134645 (*CEP192* 3), TRCN0000134623 (*CEP192* 5) (Horizon Discovery). For enforced expression of Kifc3, *KIFC3* cDNA (ENST00000541240.5) was cloned into the pGenlenti vector. For enforced expression of Cep192, human *CEP192* cDNA (clone Id: 9053008, Horizon Discovery) was inserted between attb1 and attb2 sites of the PLBC-2 lentiviral vector. Production of lentiviral supernatants by transient co-transfection of vectors with packaging plasmids (pMD2.G and pCMV-dR8.74) into 293T cells, followed by spinoculation of pre-expanded progenitors took place as per descriptions^12,19^. Post spinoculation cells underwent selection in pre-stimulation medium with 1.2 µg/ml puromycin for 3 days followed by differentiation for 5-7 days in Mk medium.

### Mice

All experiments were approved by the University of Virginia Institutional Animal Care & Use Committee (IACUC) and performed in accordance with the institutional animal care guidelines. The *Cep192^CRISPR^* strain (C57BL/6NJ-*Cep192^em1(IMPC)J^*/Mmjax), generated as part of the Knockout Mouse Phenotyping Program^20^, was purchased from MMRRC (stock #42356). The *Kifc3-/-* strain, generated by Yang et al.^21^, were purchased from MMRRC (stock #000028-MU) and backcrossed to attain >98% C57BL/6 based on Neogen miniMUGA Background Analysis V0009. PCR genotyping of genomic DNA is described in Supplement.

### Flow cytometry analysis

Human and murine progenitors cultured in Mk medium were washed, stained for surface markers -/+ DNA content, and analyzed as recently published^2,12^. For analysis of primary bone marrow Mk, murine bone marrows were harvested, processed, and labelled as described^2,12^. For all flow studies, assessments of ploidy, size (FSC), and granularity (SSC) were conducted on gated viable, singlet, CD41+ Mk; in PLBC-2 lentiviral transduction experiments the gating additionally included BFP+ cells. Samples were run on an Attune NxT flow cytometer (ThermoFisher), and analytical software consisted of FlowJo (Version 10.8.2, BD Biosciences). For the *in vitro* platelet release assay, culture-derived platelets underwent isolation and flow cytometric quantitation as described^2,12,22^. Platelet activation studies were conducted as we have published^2^, using 40 μM thrombin receptor activating peptide 6 (TRAP-6) (ThermoFisher Scientific) or 100 μM adenosine 50-diphosphate (ADP) (Sigma-Aldrich).

### Microscopy

Immunofluorescence (IF) was conducted as per previous report^2^. Cells underwent cytospin at a density of 4 x 10^4^ per glass slide followed by fixation 15 minutes in 4% paraformaldehyde/PBS at room temperature. Cells were then washed twice with PBS, and permeabilized/blocked for 1 hour in blocking buffer (PBS with 2% FBS, 2% BSA, and 0.1% Triton X-100). Centrosomal antibodies consisted of rabbit polyclonal anti-Cep170 (Proteintech, 27325-1-AP) at 1/100 or rabbit polyclonal anti-Cep192 (Proteintech, 18832-1-AP) at 1/200, combined with mouse monoclonal anti-Centrobin (Abcam, ab70448) at 1/800 in blocking buffer applied overnight at 4° C. Staining for Kifc3 employed a mouse monoclonal antibody (Santa Cruz, sc-365494) at 1/100 in blocking buffer applied overnight at 4° C. After 4 washes with blocking buffer, the secondary incubation consisted of blocking buffer with goat anti-rabbit Alexa-Fluor 488^TM^ (ThermoFisher) at 1/300, DAPI, and (where indicated) Alexa Fluor-594-Phalloidin (Invitrogen) at 1/20 for 60 minutes at room temperature. Slides washed 4 times with blocking buffer and once with PBS underwent coverslip mounting with Vectashield medium (Vector Laboratories H-1000). Images were captured with a Zeiss LSM 700 confocal microscope using the 63X oil objective and fixed signal intensity gain across compared samples. Images were processed with Fiji using maximum intensity projections of Z slices spanning entire cells. To derive a Pearson’s correlation coefficient for Cep192 and F-actin, the JACoP plugin of ImageJ was used. For quantitative comparisons, 30-60 cells were scored for each independent replicate.

### Immunoblot

For analysis of whole cell lysates, intact cell pellets underwent lysis in an equal volume per cell number of 2X Laemmli buffer (60 mM Tris-HCl, pH 6.8, 2% SDS, 100 µM dithiothreitol, 10% glycerol, and 0.01% bromophenol blue) supplemented with protease and phosphatase inhibitors (cOmplete and PhosSTOP, Roche) followed by shearing of DNA. All samples were boiled 5 minutes, followed by SDS-PAGE and immunoblot as described^23^. Primary antibodies and incubation conditions are provided in a Supplemental Table. Densitometry data were acquired and analyzed using ImageStudioLite (Version 5.2.5, LI-COR).

### In silico screening for Kifc3 inhibitors

A detailed description is provided in Supplement. We used the Enamine HTS library to identify phenyl/pyridine-based structures lacking Pan Assay Interference properties (PAINS). After generation of 3D conformations, including stereoisomers and tautomers, compounds were screened for similarity to the Kifc1 inhibitors. Virtual docking was then conducted using both the α4/α6 cleft and the α2/L5/α3 pocket of Kifc3, applying a series of programs for optimization (GOLD suite v. 5.8.1, LigandScout v. 4.4.3, PyRod, Desmond, and VMD v. 1.9.3). Candidate compounds were purchased from EnamineStore (Enamine), dissolved in DMSO, and used at the indicated concentrations. Known Kifc1 inhibitors (AZ82, CW069, and SR31527)^24–26^ were similarly acquired and employed.

### Statistics

Graphs represent results from at least three independent experiments and depict mean values ± standard error of the mean. Single, pairwise comparisons employed Student’s *t* test; multiple comparisons employed 1- or 2-way ANOVA with Tukey *post hoc* test using GraphPad Prism. A *P* value less than 0.05 was considered significant. Analysis of RNA-seq data was conducted as described^2^.

### Study approval

CD34+ cells purchased from the Fred Hutchinson Cancer Research Center, AllCells and StemCell Technologies were originally obtained from donors with informed consent and IRB approval at all institutions. Animal studies were approved and performed in compliance with the University of Virginia School of Medicine IACUC committee and guidelines.

## Results

### Control of kinesin Kifc3 levels by megakaryocyte ontogeny and Dyrk kinase activity

In a previous study, inhibitors of the kinase Dyrk1a induced an ontogenic shift in neonatal, cord blood progenitor-derived megakaryocytes, enhancing morphogenesis (enlargement and polyploidization) and platelet release to adult levels^2^. The mechanism for this shift involved the de-repression of the transcriptional co-factor Mkl1/MrtfA, known to program cytoskeletal changes associated with adult-type megakaryopoiesis^27^. To identify downstream mediators in this pathway we analyzed previously generated RNA-seq data from purified human megakaryocytes derived from: primary adult progenitors (PB), primary neonatal progenitors (CB), primary neonatal progenitors treated with a broad Dyrk inhibitor harmine (CB/H), and primary neonatal progenitors treated with a structurally distinct Dyrk1-selective inhibitor EHT 1610^28^. The only gene showing consistent, robust changes with ontogeny and with both inhibitor treatments consisted of *KIFC3*, moderately expressed in control neonatal megakaryocytes and undetectable in adult cells as well as in neonatal cells subjected to Dyrk inhibitors (Figure 1A-B). Immunoblot and immunofluorescence studies confirmed the RNA-seq findings, with Kifc3 protein levels ≥ 2-fold lower in adult megakaryocytes and in neonatal megakaryocytes treated with Dyrk inhibitors (Figures 1C-D). The immunoblot findings were verified with a second, independent antibody (Figure S1). The juxtanuclear pattern of Kifc3 seen in neonatal cells (Figure 1D) corresponds to previous descriptions of lysosomal co-localization^29^.

**Figure 1.**
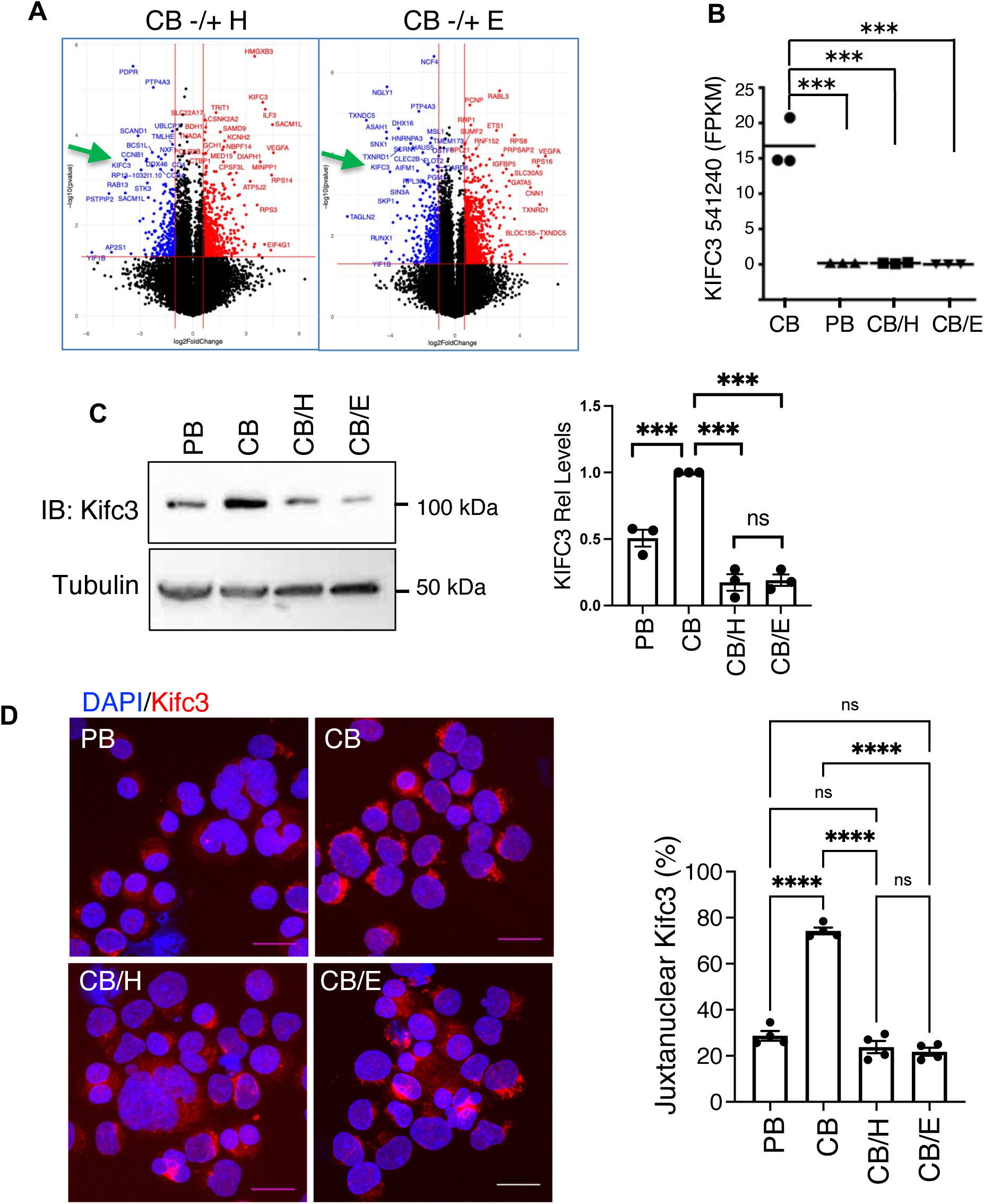
Kifc3 regulation by megakaryocyte ontogeny and Dyrk kinase activity. (A) Volcano plots from RNA-seq analysis of cord blood (CB) derived megakaryocytes treated with either DMSO, harmine (H), or EHT 1610 (E). (B) Transcript levels for full length *KIFC3* in megakaryocytes derived from adult (PB) versus cord blood progenitors -/+ Dyrk inhibitors. (C) Kifc3 protein levels in adult versus cord blood derived megakaryocytes -/+ Dyrk inhibitors. Graph depicts relative Kifc3 signal with normalization for tubulin. (D) Kifc3 localization in adult versus cord blood derived megakaryocytes -/+ Dyrk inhibitors. Scale bars = 25 µm. Graph depicts percentage of cells with discrete, juxtanuclear localization. For all graphs, each dot represents an independent experiment; ***, \*\*\*\**P* ≤ 0.005, 0.001, 1-way ANOVA with Tukey post hoc. ns = not significant. See also Figure S1.

### Implication of Kifc3 in programming of megakaryocyte ontogeny

To determine whether Kifc3 downregulation plays a role in the ontogenic shift from fetal-type to adult-type megakaryocytes, neonatal and adult progenitors underwent lentiviral shRNA knockdowns followed by megakaryocytic cultures. Two different hairpins, sh 62 and sh 63, caused mild (∼20%) or moderate (∼50%) Kifc3 decreases respectively on immunoblot (Figure S2A). In neonatal megakaryocytes, the Kifc3 knockdowns significantly enhanced polyploidization in proportion to the degree of knockdown; cells with moderate knockdown showed adult-equivalent levels of polyploidization. By contrast, Kifc3 knockdowns in adult megakaryocytes had no effect on polyploidization. (Figure 2A). Enhancement of neonatal polyploidization by Kifc3 knockdown was associated with concomitant increases in megakaryocyte size (Figure 2B) and platelet release activity (Figure 2C). The platelets produced by neonatal megakaryocytes with Kifc3 knockdown retained full responsiveness to the agonists TRAP-6 and ADP (Figure S2B). In gain-of-function experiments, enforced expression of Kifc3 in adult progenitors reproducibly suppressed megakaryocyte polyploidization and enlargement (Figure S2C), consistent with a role in determination of ontogenic phenotype.

**Figure 2.**
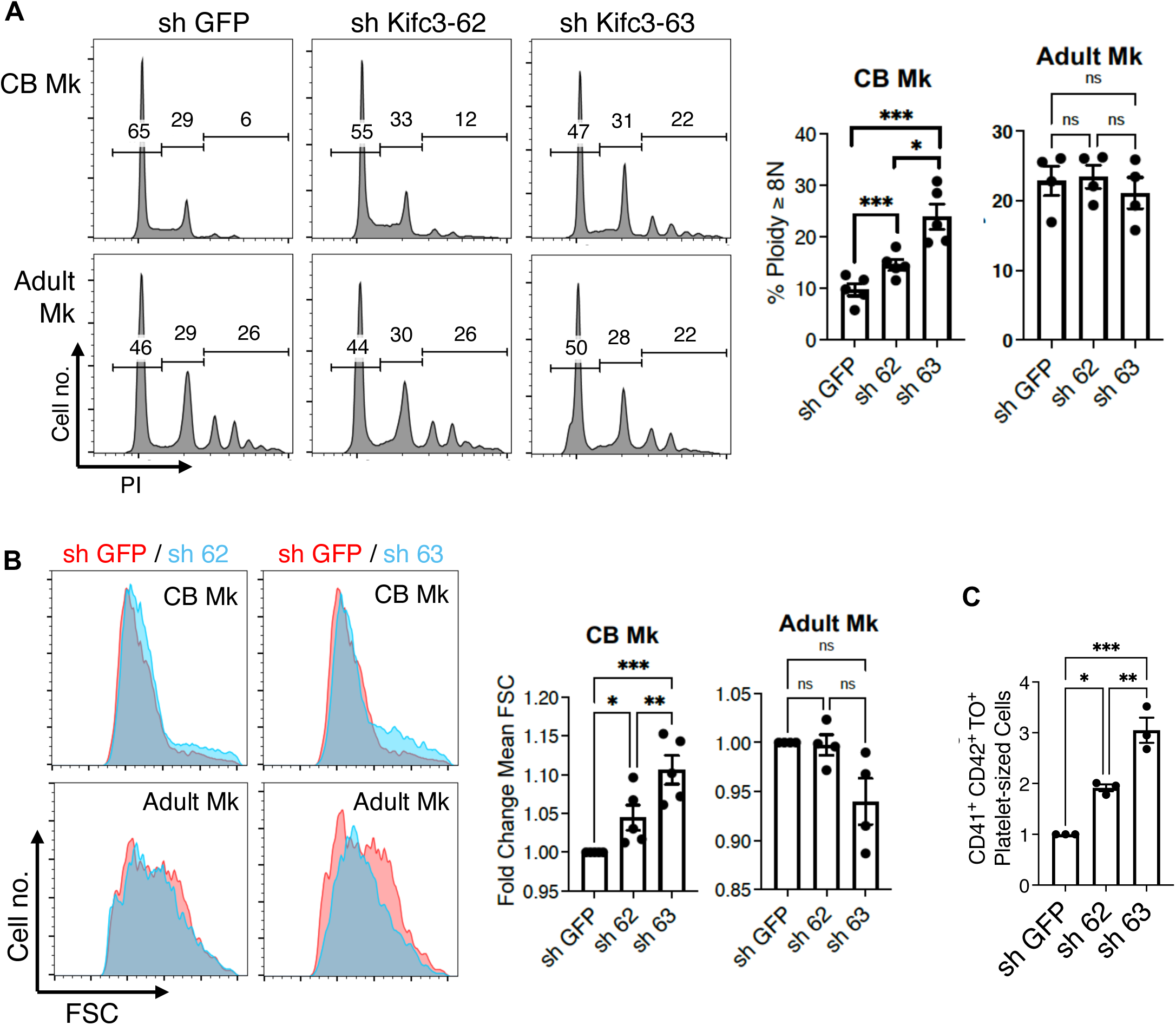
Kifc3 knockdown enhances neonatal megakaryocyte morphogenesis and thrombopoiesis. (A) Cord blood and adult progenitor derived megakaryocytes expressing control (GFP) and Kifc3 targeting short hairpin RNAs (62 and 63) were assessed for ploidy (PI) with gating on viable, singlet, CD41+ cells. Extent of knockdown is shown in Supplementary Figure 2A. (B) Assessment of cell size (FSC) in samples from Figure 2A. (C) Platelet release from neonatal megakaryocytes subjected to shRNA knockdown of Kifc3. Graph depicts relative number of platelets produced per input megakaryocyte; agonist responsiveness is shown in Supplementary Figure 2B. For all graphs, each dot represents an independent experiment; *, **, \*\*\**P* ≤ 0.05, 0.01, 0.005, 1-way ANOVA with Tukey post hoc. ns = not significant. See also Figure S2.

### The centrosomal protein Cep192 in megakaryocytes is regulated by ontogenic stage, lysosomal activity, and Kifc3

Kifc3 is a broadly expressed, minus end-directed kinesin that has been implicated in carrying several cargos, including delivery of proteins to centrosomes^30–33^. Maliga and colleagues identified Cep170 as a factor dependent on Kifc3 for its centrosomal localization in HeLa cells^31^. Immunofluorescent staining of adult megakaryocytes and neonatal megakaryocytes -/+ Kifc3 knockdown showed close co-localization of Cep170 with the daughter centriole marker centrobin, but no influence of ontogenic stage or Kifc3 levels (Figure S3). Recent studies found that a related centrosomal protein, Cep192, dynamically dissociates from centrosomes during tumor cell process extension, an activity reminiscent of adult megakaryocyte proplatelet formation^34^. Indeed, when compared with neonatal megakaryocytes, adult megakaryocytes displayed strong Cep192 upregulation and dispersion from centrosomes (Figure 3A-C); immunoblot confirmed the 3-fold increase detected on immunofluorescence (Figure 3D). Upregulation of *CEP192* transcripts were not identified in an analysis of the RNA-seq data from Figure 1A-B. Prior studies have shown tight control of Cep192 levels by ubiquitin-proteasome activity^35–37^. Treatment of neonatal and adult megakaryocytes with a panel of protease inhibitors, however, showed no effect of the proteosome inhibitors MG132 or Bortezimib (Figure S3E). Interestingly, lysosomal inhibition with either chloroquine or Bafilomycin A caused enhancement of Cep192 in neonatal but not adult megakaryocytes (Figure S3E and Figure 3E). Involvement of Kifc3 in suppression of Cep192 in neonatal megakaryocytes was demonstrated by shRNA-mediated knockdowns which induced dispersion and upregulation of Cep192, as seen in normal adult megakaryocytes (Figure 3F-I).

**Figure 3.**
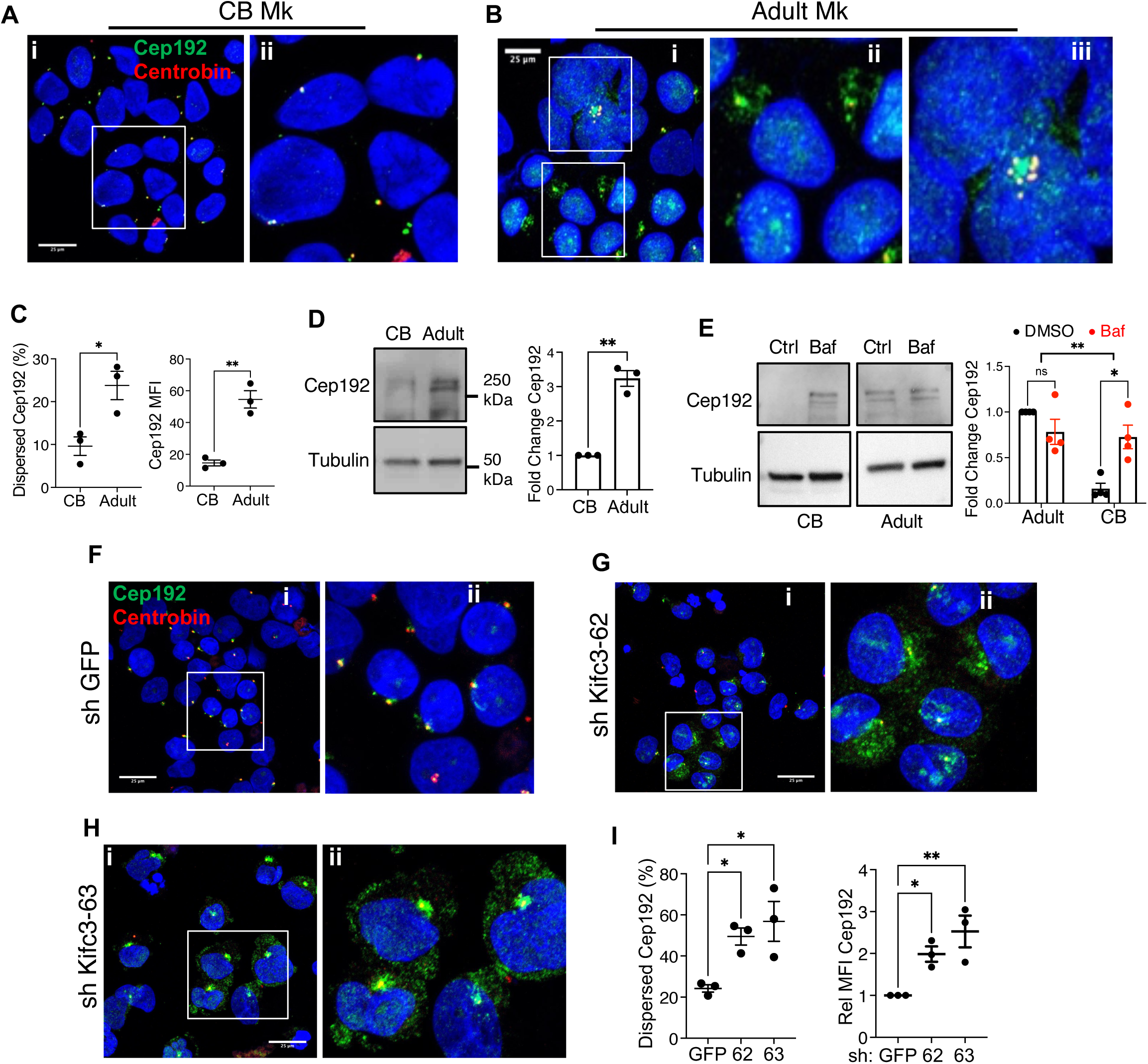
The centrosomal protein Cep192 is regulated by megakaryocyte ontogeny, lysosomal catabolism and Kifc3 levels. (A) Immunofluorescence analysis of cord blood progenitor derived megakaryocytes. ii. Expanded view of boxed region from i. (B) Analysis of adult megakaryocytes as in (A). (C) Quantitation of immunofluorescence results, showing percentage of cells with dispersed Cep192 and mean fluorescence intensity of Cep192 signal in arbitrary units. (D) Cep192 levels by immunoblot, with graph showing tubulin-normalized relative levels. (E) Neonatal and adult progenitors cultured 6 days in megakaryocyte medium with 10 nM Bafilomycin A (red dots) for the last 24 hours. Images cropped from same gel/exposure. (F-I) Immunofluorescence analysis of cord blood progenitor derived megakaryocytes subjected to short hairpin RNAs (GFP, 62, 63). Scale bars = 25 µm. (H) Percentage of cells with dispersed Cep192 and mean fluorescence intensity of Cep192. Each dot represents an independent experiment; *, \*\**P* ≤ 0.05, 0.01, Student’s *t* test (C-D), 2-way ANOVA (E) or 1-way ANOVA with Tukey post hoc (I). See also Figure S3.

### Cep192 is a mediator of adult megakaryocyte morphogenesis

To assess the involvement of Cep192 in adult megakaryocyte morphogenesis, adult progenitors underwent lentiviral shRNA knockdowns followed by megakaryocytic cultures. Three different hairpins, 1, 3 and 5, diminished Cep192 on immunoblot to levels approximating those in neonatal megakaryocytes (Figure S4A). In all cases, Cep192 knockdown significantly diminished polyploidization and enlargement (Figure 4A). Because Cep192 participates in centrosome biogenesis, the effects of its knockdown might reflect a role of centrosome biogenesis in adult megakaryocyte morphogenesis. However, treatment of adult progenitors with centrinone, an inhibitor of centrosome biogenesis, enhanced megakaryocyte polyploidization (Figure S4B). The contribution of Cep192 to megakaryocyte morphogenesis is thus likely to be independent of its role in centrosome biogenesis. For *in vivo* studies, megakaryocytes from *Cep192* mutant mice were analyzed. The mutation consisted of a CRISPR-mediated deletion of exon 5, which prevented protein expression for 5 out of 7 isoforms, but retained expression two forms with amino terminal truncations of 230 and 632 amino acids (Figure S4C). Marrow megakaryocytes from adult homozygous mutant mice manifested polyploidization deficiencies in both males and females, when compared with wild type littermates (Figure 4B). In gain-of-function experiments, we transduced human neonatal progenitors with pLBC-2 lentiviral expression constructs carrying a BFP marker. In these experiments, enforced expression of Cep192 in neonatal Mk strongly enhanced morphogenesis, with a consistent 2-fold increase in polyploidization, as well as a significant increase in size (Fig. 4C).

**Figure 4.**
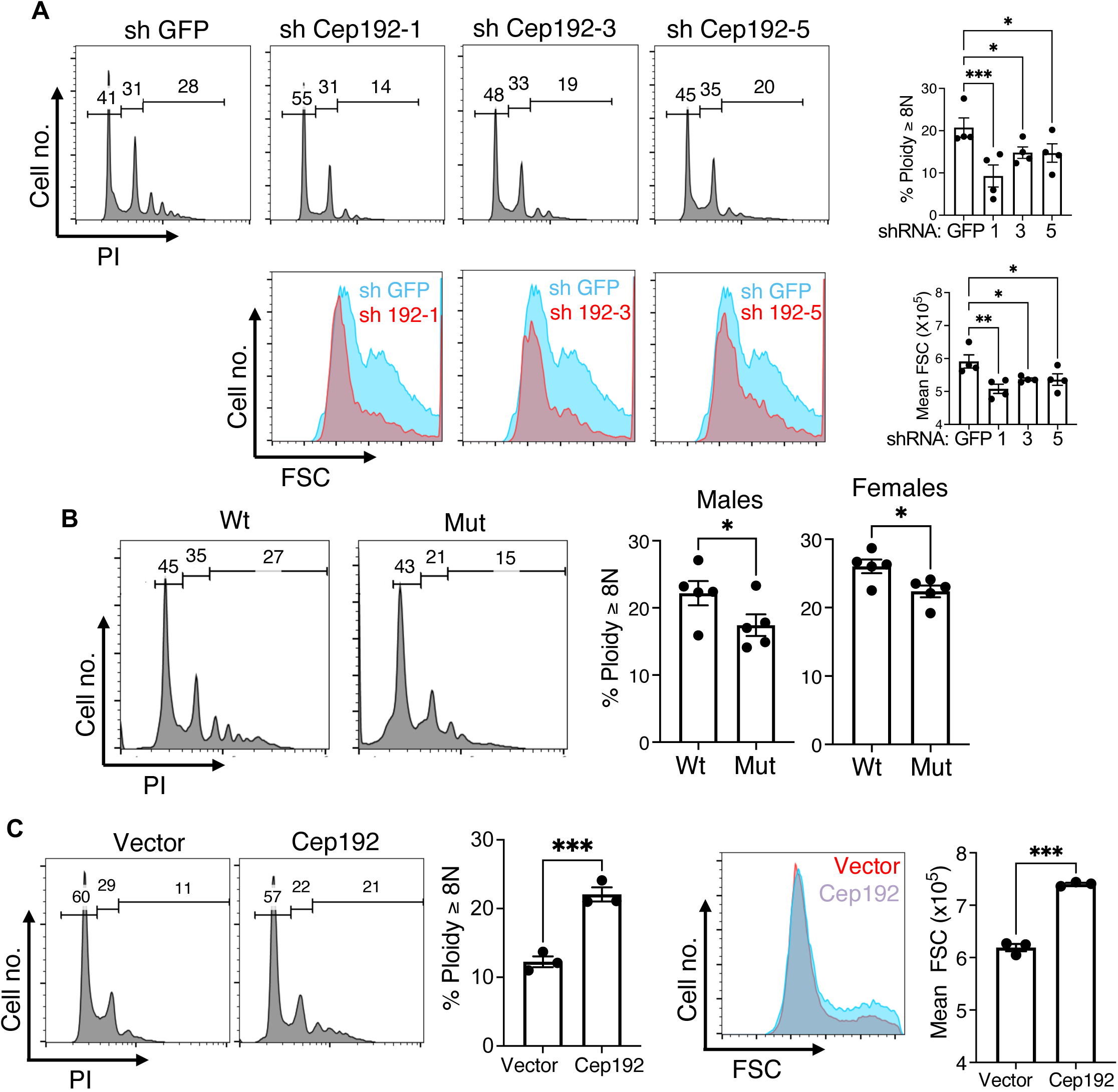
Cep192 contributes to adult megakaryocyte morphogenesis. (A) Adult progenitor derived megakaryocytes expressing control (GFP) and Cep192 targeting short hairpin RNAs (1, 3, and 5) were assessed for ploidy (PI) and size (FSC) with gating on viable, singlet, CD41+ cells. Extent of knockdown is shown in Supplementary Figure 4A. (B) Marrow from adult wild type (Wt) and homozygous *Cep192* mutant (Mut) mice was analyzed for megakaryocyte ploidy with gating on viable, singlet, CD41+ cells. Graph summarizes findings for males (left) and females (right). Diagram of *Cep192* locus mutation is shown in Supplementary Figure 4C. (C) Neonatal progenitor derived megakaryocytes transduced with control or Cep192 expression vector were assessed for ploidy (PI) and size (FSC) with gating on viable, BFP+, singlet, CD41+ cells. For all graphs, each dot represents an independent experiment or subject; *, **, \*\*\**P* ≤ 0.05, 0.01, 0.005, 1-way ANOVA with Tukey post hoc for (A) and Student’s *t* test for (B). See also Figure S4.

### Boosting morphogenesis and platelet production in neonatal megakaryocytes through small molecule targeting of Kifc3

Kinesin motor domains are amenable to small molecule inhibition for therapeutic purposes^38^. Specific, bioactive inhibitors of Kifc1, a Kifc3 paralog, have been identified by multiple approaches including high throughput drug screening and computational modeling^17,24–26^. To identify Kifc3 inhibitors, we employed a strategy of computer aided drug design, using the workflow shown in Figure 5A. Candidate compounds, as well as known Kifc1 inhibitors, were screened for effects on neonatal megakaryocyte morphogenesis resembling Kifc3 knockdown. Notably, none of the Kifc1 inhibitors (AZ82, SR31527, and CW069) used at known bioactive doses^24–26^ had any effects in this assay (Figure S5A-B). One of the initial candidate compounds, LS5, caused small but consistent increases in ploidy and size at 40 µM (Figure S5C-D). LS5 was predicted to bind the α2/L5/α3 pocket of the Kifc3 motor domain (Figure 5B-C). Secondary screening of 14 LS5 analogs identified one compound, A1, with significantly enhanced bioactivity (Figure S5C-D). A1 structurally differed from LS5 minimally but exerted robust effects on neonatal megakaryocyte morphogenesis at 20 µM, akin to those seen with moderate Kifc3 knockdown (Figure 5B, D-E). Importantly, A1 significantly enhanced platelet release from neonatal megakaryocytes (Figure 5F). A1 had no effects on other neonatal lineages except to slightly enhance erythroid viability (Figure S5E-F), suggesting megakaryocyte specificity. To determine whether A1 exerted its effects through targeting of Kifc3, fetal liver progenitors from *Kifc3+/-* and *-/-* embryos underwent megakaryocytic culture -/+ 20 µM A1. Differential drug responsiveness based on genotype supported Kifc3 as a key target for A1 activity (Figure 5G).

**Figure 5.**
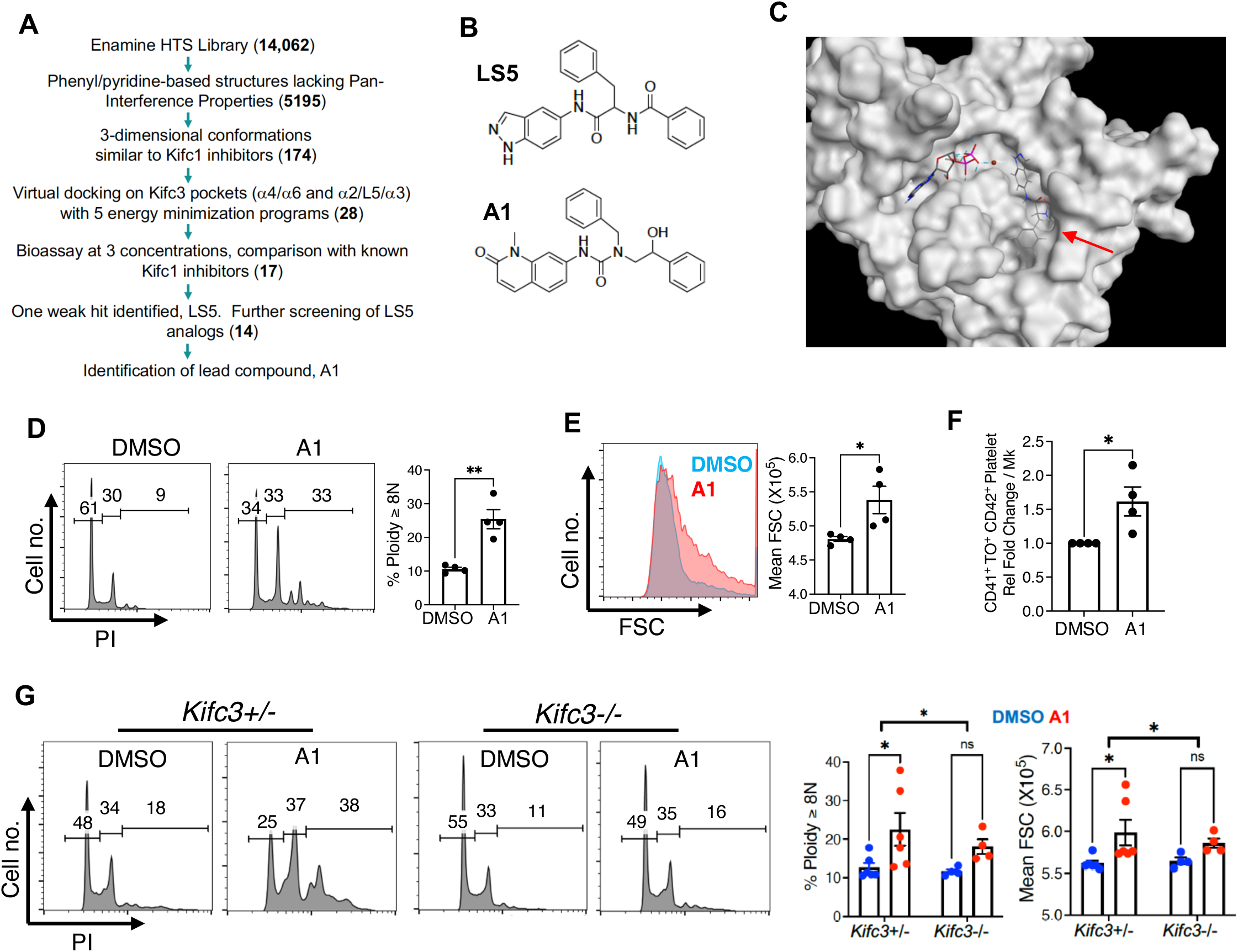
Small molecule targeting of Kifc3 to enhance neonatal morphogenesis and thrombopoiesis. (A) Workflow for *in silico* screening, with compound numbers at each step shown in parentheses. (B) Structures of initial hit LS5 and lead compound A1. (C) Model of LS5 (red arrow) docking in the α2/L5/α3 pocket of the Kifc3 motor domain. Also shown are ADP with Mg^2+^ ion (red ball). (D-E) Cord progenitors subjected to megakaryocytic cultures were treated with 20 µM A1 or DMSO followed by analysis of analysis of ploidy (D) and size (E) with gating on viable, singlet CD41+ cells. (F) Platelet release from neonatal megakaryocytes treated with 20 µM A1 or DMSO. (G) Fetal liver progenitors from *Kifc3+/-* and *Kifc3-/-* embryos were treated with 20 µM A1 or DMSO followed by analysis of ploidy and size, with gating on viable, singlet CD41+ cells. For all graphs, each dot represents an independent experiment or subject; *, \*\**P* ≤ 0.05, 0.01, Student’s *t* test for (D-F) and 2-way ANOVA with comparison of A1 responsiveness by genotype for (G). See also Figure S5.

### Small molecule targeting of Kifc3 induces molecular features of adult megakaryopoiesis

Adult and neonatal megakaryocytes possess distinct expression patterns for Cep192 (see Figure 3A-B) and F-actin^2^. The ability of Cep192 to promote actin remodeling^34^ suggests an inter-relationship between these two findings. As seen with Kifc3 knockdown, A1 treatment of neonatal progenitors caused an increase and dispersion of Cep192 in megakaryocytes (Figure 6A-B). Immunofluorescence demonstrated minimal co-localization of Cep192 with F-actin in neonatal megakaryocytes, in contrast to the high co-localization in adult megakaryocytes. Notably, A1 treatment significantly augmented this co-localization in neonatal megakaryocytes, recapitulating the sheets of cytoplasmic F-actin forming at the sites of cytoplasmic Cep192 clouds (Figure 7A).

**Figure 6.**
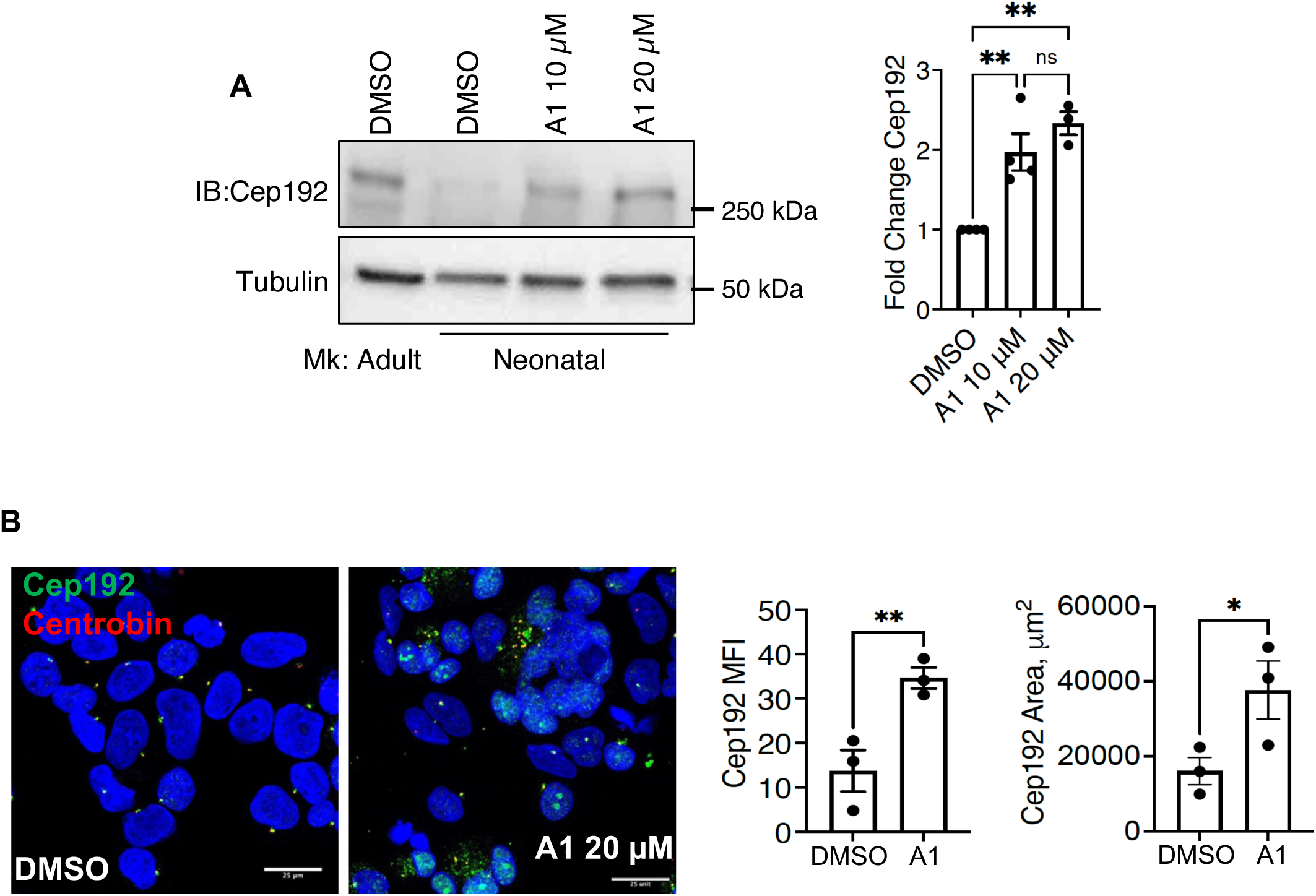
Small molecule targeting of Kifc3 in neonatal megakaryocytes induces Cep192 upregulation and dispersion. (A) Immunoblot levels of Cep192. Graph depicts tubulin-normalized results from neonatal megakaryocytes treated with DMSO or the indicated doses of A1. (B) Immunofluorescence analysis of neonatal megakaryocytes treated with DMSO or 20 µM A1. Scale bars = 25 µm. Graphs show mean fluorescence intensity in arbitrary units and degree of dispersion reflected by signal area. For all graphs, each dot represents an independent experiment; *, \*\**P* ≤ 0.05, 0.01, ANOVA with Tukey post hoc for (A) and Student’s *t* test for (B).

**Figure 7.**
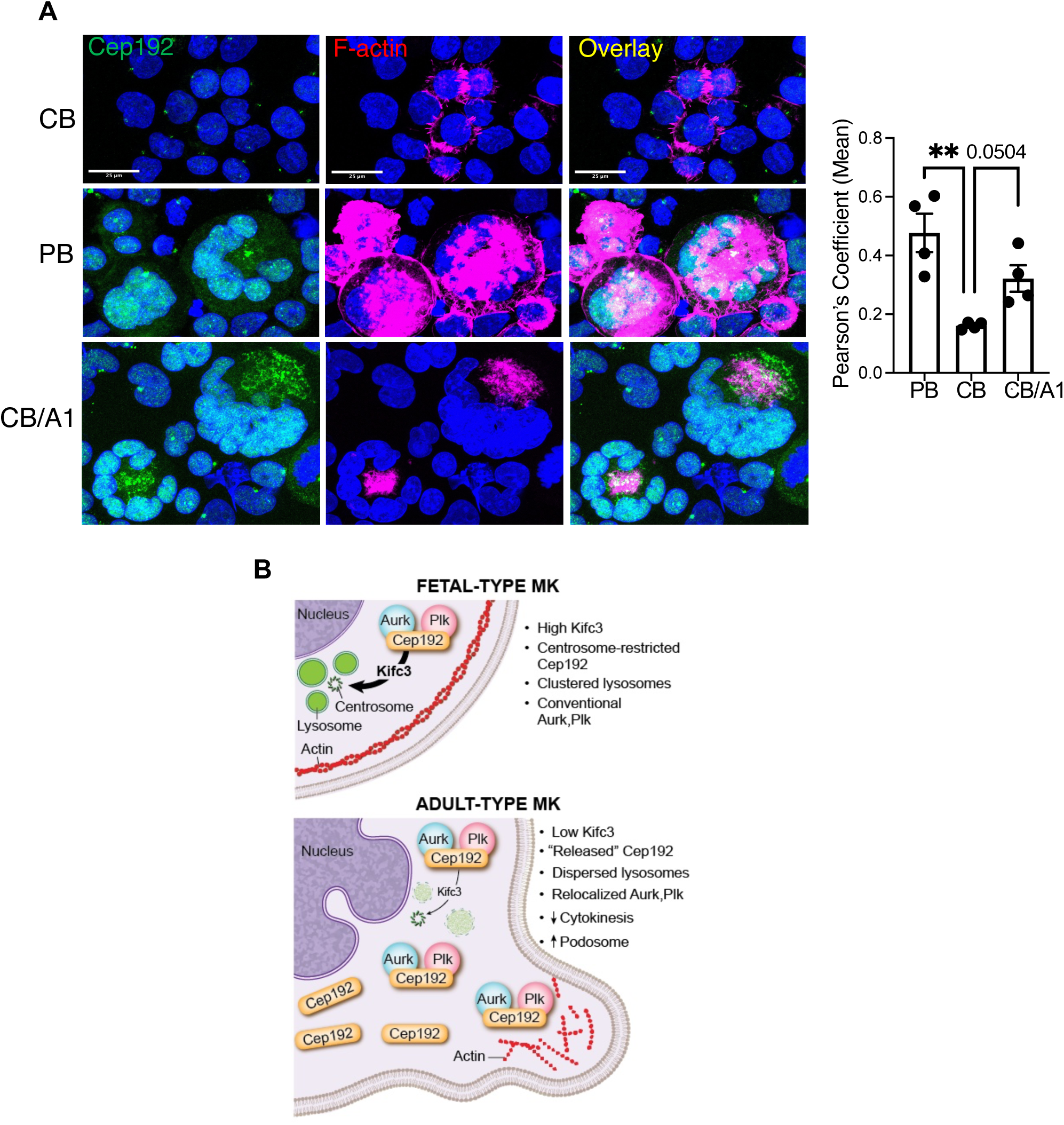
F-actin colocalization with Cep192 is influenced by megakaryocyte ontogeny and small molecule targeting of Kifc3. (A) Immunofluorescence analysis of adult (PB) megakaryocytes or neonatal megakaryocytes (CB) -/+ treatment with 20 µM A1. Scale bars = 25 µm. Graph depicts the colocalization index for the Cep192 and F-actin signals. (B) Diagram of proposed mechanism for Kifc3 regulation of megakaryocyte ontogeny. For all graphs, each dot represents an independent experiment; \*\**P* ≤ 0.01, ANOVA with Dunnett post hoc.

## Discussion

The capacity to control ontogenic phenotype has therapeutic importance in multiple settings. Reverting erythropoiesis to a fetal mode offers an effective treatment strategy for anemias with β-globin defects, such as sickle cell disease and thalassemia. Driving fetal-to-adult switching in megakaryocytes has potential to ameliorate thrombocytopenias associated with premature infancy and umbilical cord blood stem cell transplantation. In addition, megakaryocyte neoplasms in neonates (associated with t(1;22) or trisomy 21) employ oncofetal drivers and may be treatable with ontogenic switching agents. Our previous study showed that Dyrk1a inhibitors effectively induce fetal-to-adult switching in human megakaryocytes from several sources: cord blood progenitors, SR1-expanded cord blood progenitors, iPSC, and immortalized iPSC^2^. This approach has translational potential for *ex vivo* platelet production but is not applicable for patient treatment due to the broad toxicity associated with the Dyrk inhibition. The current studies thus sought to identify the downstream components of the Dyrk1a-Mkl1/MrtfA ontogenic pathway.

Kifc3 fulfills several criteria as a downstream mediator of this pathway. Firstly, its expression differed between neonatal and adult megakaryocytes, and treatment of neonatal cells with Dyrk inhibitors shifted levels downward to those seen in adult cells. The mechanism for ontogenic regulation of *Kifc3* likely involves either direct or indirect repression by Mkl1/MrtfA complexes. In muscle differentiation, which has several parallels with megakaryopoiesis^39^, Mkl1/MrtfA represses MyoD^40^ which in turn promotes expression of *Kifc3*^41^. Secondly, shRNA-mediated downmodulation of Kifc3 in neonatal megakaryocytes strongly enhanced morphogenesis and platelet production, while enforced expression in adult megakaryocytes impaired morphogenesis. Thirdly, Kifc3 antagonism in neonatal cells elicited molecular-cytoskeletal features specific to adult megakaryocytes: dispersion and upregulation of Cep192, as well as formation of sheets of F-actin.

A potential mechanism for Kifc3 control of megakaryocyte ontogeny is shown in Figure 7B. As a minus end-directed kinesin, Kifc3 transports a variety of cargos toward the centrosome. Notable cargos include components of the centrosome as well as lysosomes^29,31,33^. With regard to the latter, Bejarno et al. identified a role for Kifc3 in juxtanuclear clustering of lysosomes and showed impairment in autophagic flux associated with *Kifc3* knockout^29^. Of relevance, our results implicate a lysosomal mechanism in the ontogenic regulation of Cep192 (Figure 3E) and suggest that Kifc3 in neonatal megakaryocytes functions to coordinate efficient delivery of Cep192 to lysosomes. Whether Cep192 is a direct cargo of Kifc3 or is affected in a secondary manner by peri-centrosomal clustering of lysosomes remains unknown. Prior studies have shown only proteosomal catabolism of Cep192^35–37^, but precedents exist for specialized autophagy of centrosomal elements^42–44^.

Cep192 fulfills multiple criteria as an ontogenic effector downstream of Kifc3. Firstly, Cep192 levels and localization differed between neonatal and adult megakaryocytes. Secondly, Cep192 changed from a fetal to adult pattern in neonatal megakaryocytes subjected to Kifc3 knockdown or inhibition. Thirdly, knockdown of Cep192 in adult progenitors impaired megakaryocyte morphogenesis. This effect most likely did not result from impaired centrosome biogenesis because the degree of knockdown was modest (∼50%) and because treatment of adult megakaryocytes with a known inhibitor of centrosome biogenesis^45^ actually enhanced morphogenesis. Fourthly, mice homozygous for *Cep192* mutation displayed defects in megakaryocyte polyploidization. This mutation is predicted to eliminate Cep192 isoforms with the amino terminal Plk4 binding site, suggesting or role for this interaction in megakaryopoiesis. The mechanism for Cep192 control of morphogenesis likely relates to a novel function described by Luo et al. in which its translocation from the centrosome to the cell periphery promotes actin remodeling and process extension^34^. This activity is mediated by its recruitment of the kinases Plk4 and Aurkb which act in a semi-redundant fashion. Our data show co-localization of Cep192 with F-actin specifically in adult megakaryocytes and in neonatal megakaryocyte treated with Kifc3 inhibitor. We therefore propose that Cep192 coordinates adult megakaryocyte morphogenesis through the shifting of kinases away from the centrosome-associated mitotic apparatus, resulting in endomitosis, and redirecting them to the peripheral actin cytoskeleton, promoting cytoplasmic enlargement and process extension.

Kifc3 represents a desirable target for therapeutic manipulation of megakaryocyte ontogeny. The fetal-adult shift occurs through its loss of function. Its kinesin motor domain has been structurally characterized by X-ray crystallography. Crystal structures of the closely related Kifc1 motor domain in complex with specific inhibitory compounds have also been solved^17^. *Kifc3* knockout mice are viable and fertile, with no significant abnormalities^21^. We followed an *in silico* approach, similar to one used to identify a Kifc1 inhibitor^25^, followed by bioassay screening to obtain a lead compound A1 that extensively recapitulates the effects of Kifc3 knockdown. A1 likely exerts its effects through Kifc3 inhibition because *Kifc3*-/- megakaryocytes fail to respond, and Kifc1 inhibitors have no effect in the bioassay. Using in vitro enzymatic assays with purified recombinant Kifc3 motor domain fused to GST, however, failed to demonstrate inhibitory activity by A1 (not shown). The basis for the discrepancy between *in vivo* and *in vitro* findings is unclear. Nevertheless, these studies provide an approach for developing therapeutic ontogenic switching agents that act on a newly established molecular pathway.

## Acknowledgments

We thank the UVA Flow Cytometry Core Facility for assistance with flow cytometry; the UVA Advanced Microscopy Facility for assistance with confocal imaging; Jadranka Loncarek and Anita Eckly-Michel for helpful suggestions; Bon Trinh for reviewing the manuscript. This work was supported by grants from the National Institutes of Health (R01 HL149667, R01 DK079924, R56 DK141123).

## Author Contributions

K.E.E. performed experiments, conceptualized and interpreted data, and wrote the manuscript. S.L. conducted *in silico* screening for Kifc3 inhibitors and produced molecular models of compounds binding to the motor domain. C.W. and G.W. provided supervision for the *in silico* screening and the molecular modeling. V.B., R.S., and D.M.K. performed experiments and interpreted data. A.N.G. supervised the project, interpreted the data, and wrote the manuscript.

