## Supplement for "The Kifc3 Motor Protein Controls Centrosomal Factor Cep192 in Ontogenic Coordination of Megakaryocyte Development"

**Supplementary Figure 1**

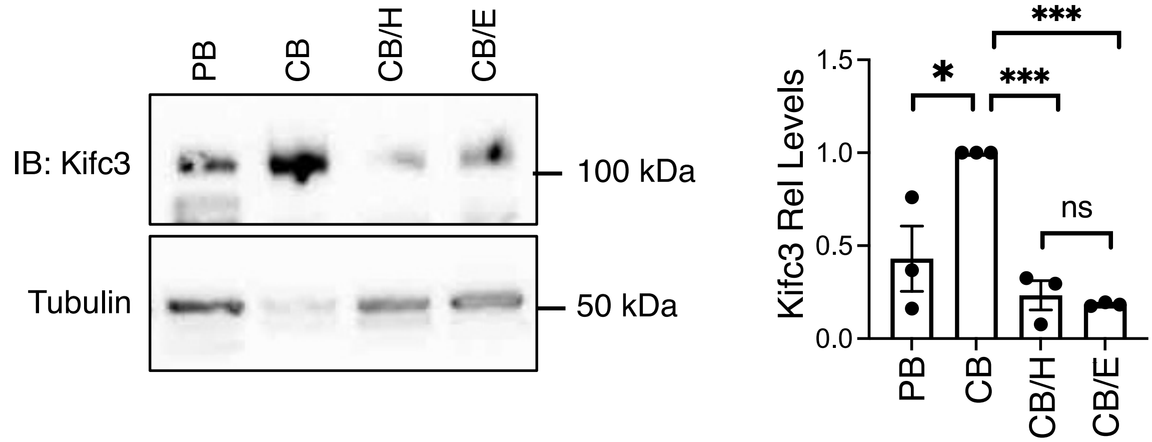


**Supplementary Figure 1. Kifc3 regulation by megakaryocyte ontogeny and Dyrk kinase activity, confirmation with an independent antibody.** Kifc3 protein levels in adult versus cord blood derived megakaryocytes -/+ Dyrk inhibitors. Graph depicts relative Kifc3 signal with normalization for tubulin. These experiments used a mouse monoclonal antibody from Santa Cruz instead of the rabbit polyclonal antibody from Sigma used in Figure 1C. Each dot represents an independent experiment; $^{*,****}P \leq 0.05$, 0.001, 1-way ANOVA with Tukey post hoc. ns = not significant.

**Supplementary Figure 2**

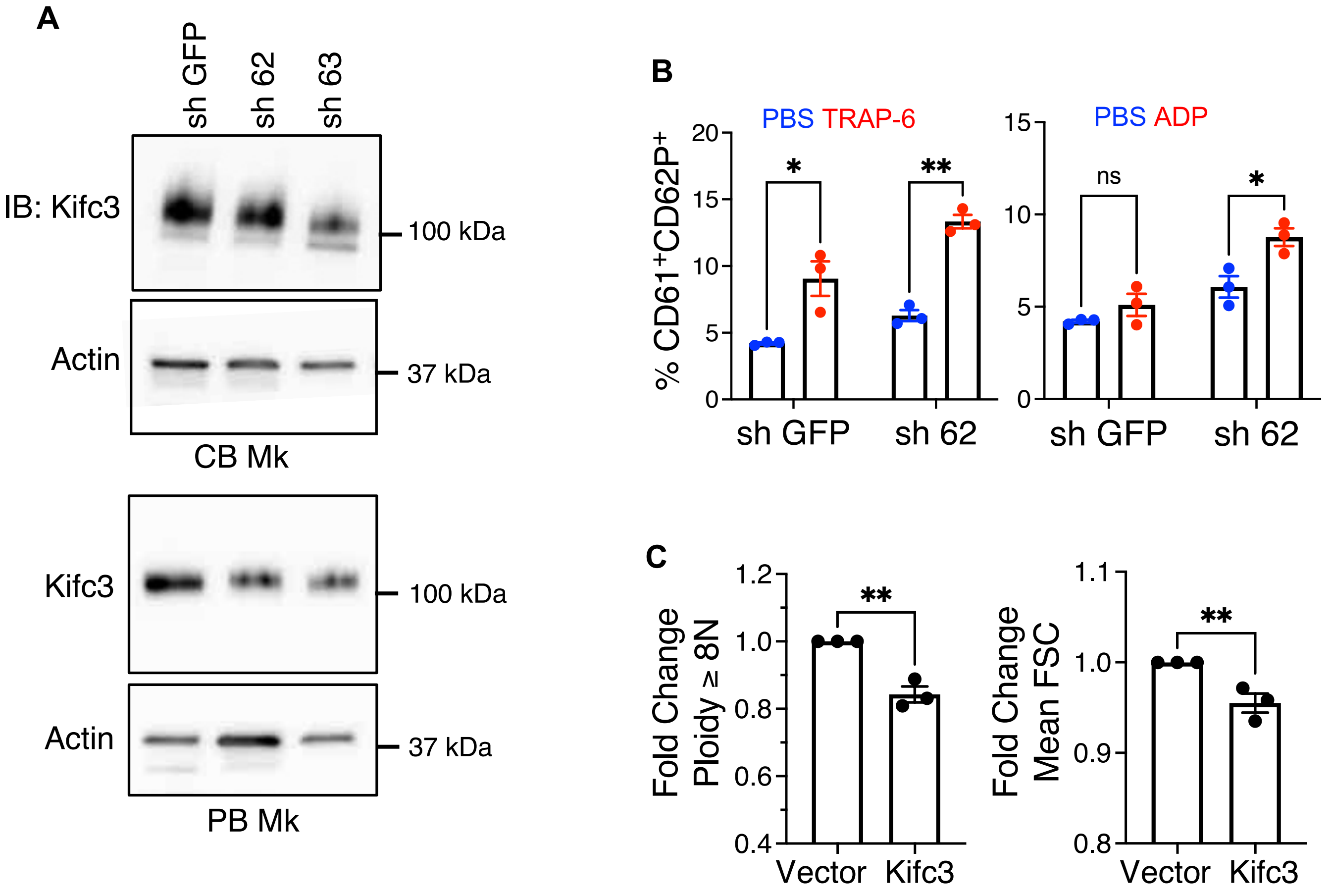


**Supplementary Figure 2. Kifc3 manipulation in human progenitor derived megakaryocytes.** (A) Kifc3 protein levels in adult versus cord blood derived megakaryocytes subjected to control or Kifc3 targeting shRNA expression. (B) Flow cytometric analysis of platelet upregulation of CD62P in response to agonists. (C) Adult progenitors transduced with lentiviral parent vector or expression construct for Kifc3 were assessed for ploidy and size (FSC) with gating on viable, singlet CD41+ cells. Each dot represents an independent experiment; *,***P* ≤ 0.05, 0.01, Student's *t* test. ns = not significant.

**Supplementary Figure 3**

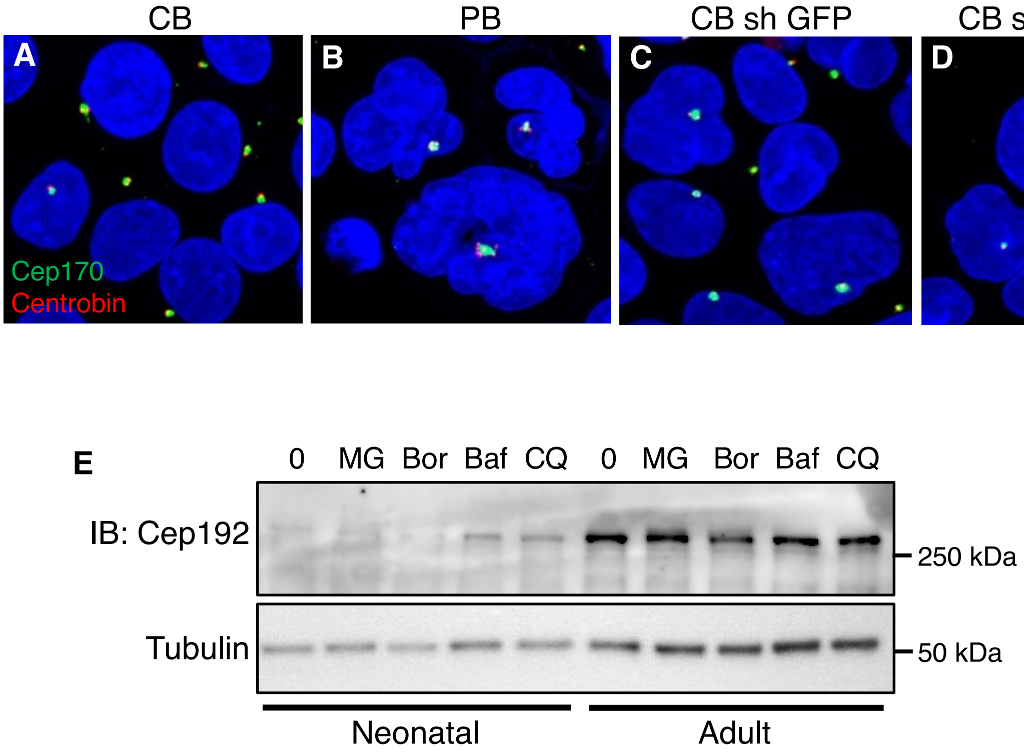


**Supplementary Figure 3. The centrosomal protein Cep170 is not affected by megakaryocyte ontogeny or Kifc3 levels; lysosomal regulation of Cep192 in neonatal but not adult megakaryocytes.** (A-B) Immunofluorescence analysis of megakaryocytes derived from cord blood (A) or adult (B) human progenitors. (C-D) Immunofluorescence analysis of megakaryocytes derived from cord blood subjected to control (C, GFP) or Kifc3 targeting (D, 63) short hairpin RNA expression. (E) Neonatal and adult progenitors underwent megakaryocytic cultures for 6 days with the following inhibitors added for the last 6 hours: MG132 (MG) at 20 μM, Bortezimib (Bor) at 0.5 μM, Bafilomycin A (Baf) at 0.5 μM, and chloroquine (CQ) at 40 μM. Whole cell lysates were subjected to immunoblot.

**Supplementary Figure 4**

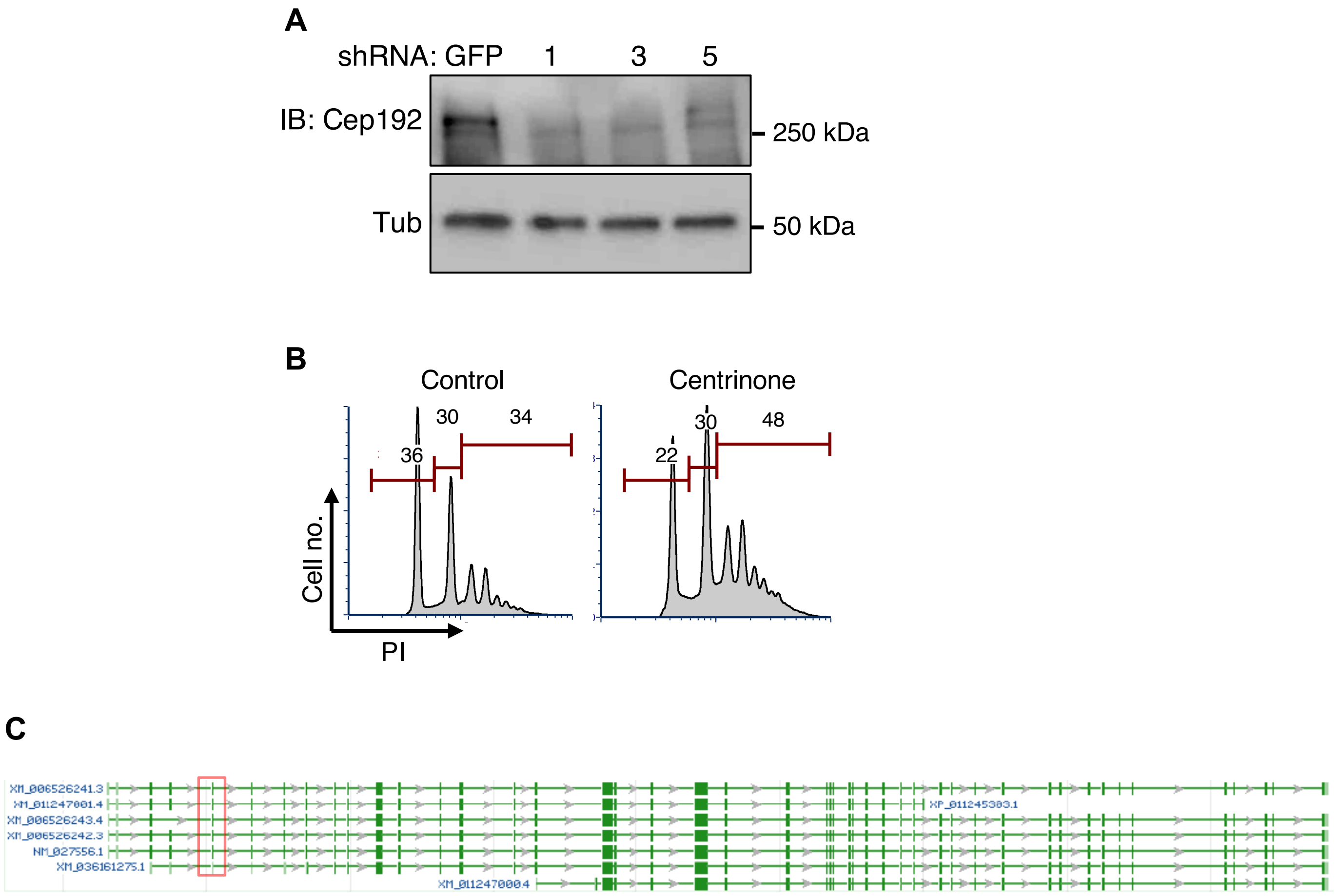


**Supplementary Figure 4. Cep192 in megakaryocyte morphogenesis.** (A) Cep192 protein levels in adult derived megakaryocytes subjected to control or Cep192 targeting shRNA expression. (B) Ploidy analysis of adult human megakaryocytes subjected to blockade of centrosome biogenesis by treatment with the Plk4 inhibitor centrinone. (C) Diagram of murine *Cep192* genomic locus illustrating alternatively spliced isoforms. Red box shows exon 5, which is deleted by CRISPR mutagenesis.

**Supplementary Figure 5**

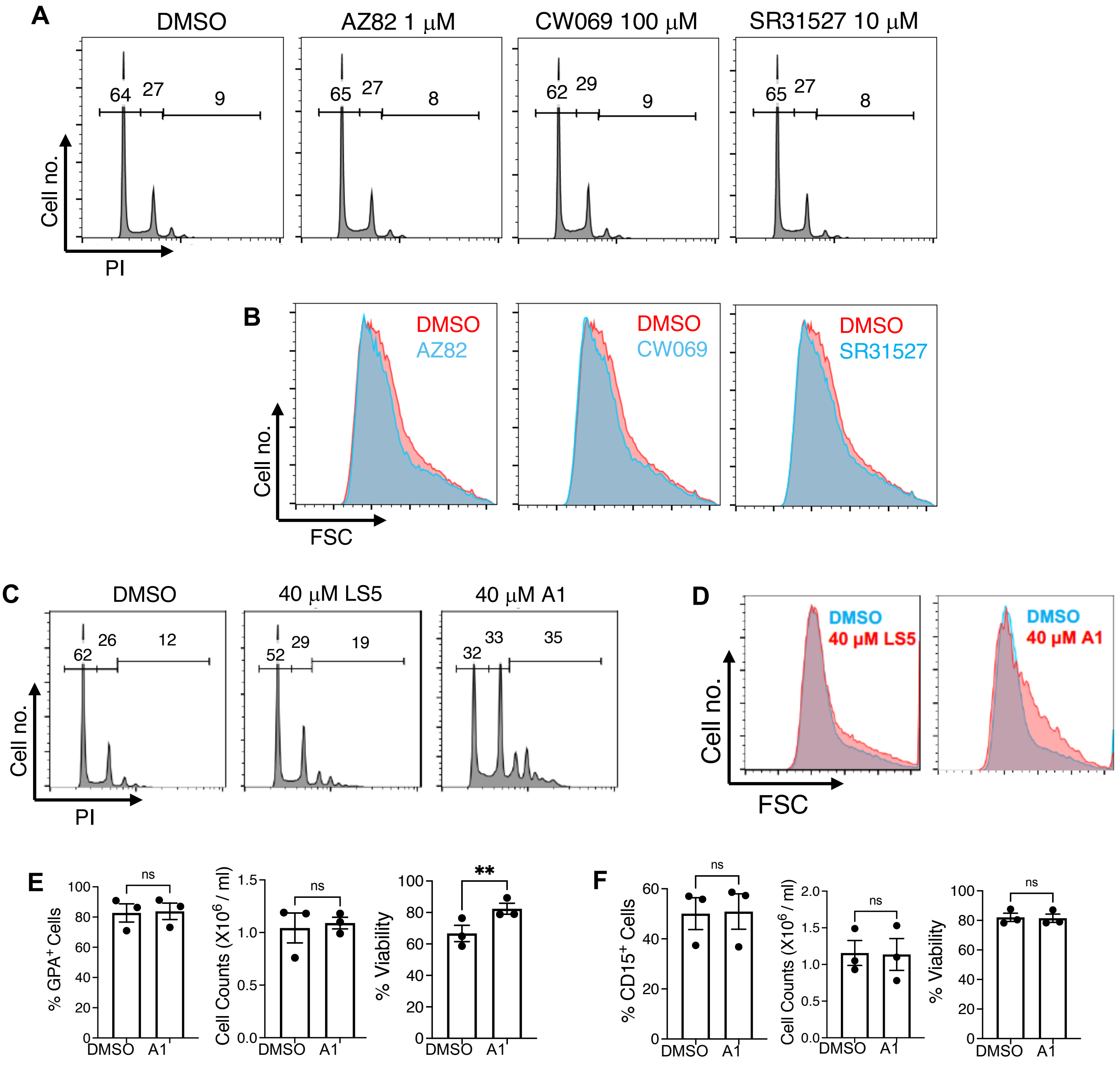


**Supplementary Figure 5. Small molecule targeting of Kifc3 enhances megakaryocyte morphogenesis and does not affect other lineages.** (A-B) Cord blood progenitors subjected to megakaryocytic cultures were treated with Kifc1 inhibitors (AZ82, CW069, SR31527) or DMSO followed by analysis of analysis of ploidy and size with gating on viable, singlet CD41+ cells. (C-D) Cord blood progenitors subjected to megakaryocytic cultures were treated with LS5, A1 or DMSO followed by analysis of analysis of ploidy and size with gating on viable, singlet CD41+ cells. (E-F) Cord blood progenitors subjected to erythroid (E) or granulocytic (F) cultures with 20 μM A1 or DMSO were analyzed for differentiation, cell counts, and viability. Each dot represents an independent experiment; *,***P* ≤ 0.05, 0.01, Student's *t* test. ns = not significant.

**Mice genotyping**

The genotypes of *Cep192*$^{CRISPR}$ strain knockout mice were determined by PCR using wild type forward primer (5′ TTG CTT GGT ATC TGT CTT CTG C 3′), common reverse primer (5′ CAT GGT TCC CAA GAC AAA GG 3′) and mutant forward primer (5’ CAT GAG AGG CAG GGT GAT G 3’). PCR reactions were run in 3% agarose gel for 3 hours. The PCR generates 108-bp wild-type and 116-bp mutant bands. For genotyping *Kifc3-/-* strain PCR were performed with four primers, P1 (5’ TGCAGCGGCAGGTGCTGAAG 3’), P2 (5’AGGTTCTCGTGTACTGCCTT 3’), P3 (5’ CAGTCAACAGCAACTGATGG 3’), P4 (5’ GGACAGGTCGGTCTTGACAA 3’). The PCR generates 800-bp wild-type and 400-bp mutant bands. PCR protocols for genotyping are described in MMRRC website.

**Antibodies Used for Immunoblots:**

| Antibody | Species | Company (Catalog number) | Dilution (Buffer) | Duration (temperature) |
|---|---|---|---|---|
| Anti-Kifc3 | Mouse | Santa Cruz (clone D-9. SC-365494) | 1:500 (1% milk in *TBST) | Overnight (4° C) |
| Anti-Kifc3 | Rabbit | Sigma (Protein Atlas. HPA021240) | 1:1000 (1% milk in TBST) | Overnight (4° C) |
| Anti-Cep192 | Rabbit | Fortis Life Sciences (Bethyl-A302-324A) | 1:2000 (2% milk in TBST) | Overnight (4° C) |

*TBST: Tris buffer solution plus 0.1% Tween 20 (pH 7.5).
All membranes were blocked with 5% milk in TBST for one hour at room temperature.
Secondary antibodies were diluted in 1% milk in TBST 1:10,000.

### *In silico* screening

Database preparation

Enamine HTS database was selected as the starting point, since it contains accessible ligands and smaller ones that have capacity to be optimized. Substructure of phenyl/pyridine-alanine was filtered as aim molecule. PAINs[1] were filtered out with programed substructure filtering tools supplied by RDKit. The molecules were protonated by 'protonation 3D' function implanted in MOE v.2020.0901 (Molecular Operating Environment; Chemical Computing Group ULC, Montreal, QC, Canada). Molecules' 3D structures were generated by Idbgen in LigandScout v.4.4.3 [10, 11] and Oeomega Classic in Openeye. KIFC1 inhibitors were searched with ROCS, Openeye, with TanimotoCombo scoring function.

Molecular Docking

Selected molecules were docked into the Kifc3 motor domain protein structure with GOLD suite v. 5.8.1. Search efficiency was set to 200% and the genetic algorithm (GA) was set to run 10 times for each ligand. The docking center was defined by the position of Phe626 para-C atom and a sphere of 10 Å around it. The resulting docking poses were energy minimized employing the MMFF94 force field implemented in LigandScout v. 4.4.3, in which, subsequently, visual inspection and 3D pharmacophore modeling was done.

Trace Water-protein interactions during molecular dynamics (MD) simulations

We used PyRod, a free and open-source python software designed by our group, to trace the thermodynamic properties of water molecules in the apo Kifc3 protein α2/L5/α3 or α4/α6 binding site during MD simulation, based on the idea that ligands entering a protein binding pocket essentially compete with water molecules for binding to the protein[2]. All-atom MD simulations were performed with the prepared model of Kifc3 motor domain as required by PyRod's input. The protein was solvated in a 15 Å padding box with TIP4P water model. Sodium ions were added to the environment system for electric neutrality. A default seven-step protocol for system relaxation and equilibration was implemented. The temperature and pressure for simulation were set at 300 K and 1.01325 bar, respectively. Prepared MD simulation systems were carried out with the Desmond[3] on water-cooled Nvidia RTX 2080 Ti graphical processing units (GPU) for 10 ns in ten replicas using the OPLS-AA force field, with recorded interval of 10 ps. Trajectories were wrapped and aligned in VMD v. 1.9.3[4].

1. Baell JB, Holloway GA. New Substructure Filters for Removal of Pan Assay Interference Compounds (PAINS) from Screening Libraries and for Their Exclusion in Bioassays. *Journal of Medicinal Chemistry*. 2010;53(7):2719-2740.
2. Schaller D, Pach S, Wolber G. PyRod: Tracing Water Molecules in Molecular Dynamics Simulations. *Journal of Chemical Information and Modeling*. 2019;59(6):2818-2829.
3. Bowers KJ, Chow DE, Xu H, et al. Scalable Algorithms for Molecular Dynamics Simulations on Commodity Clusters. SC '06: Proceedings of the 2006 ACM/IEEE Conference on Supercomputing; 2006:43-43.
4. Humphrey W, Dalke A, Schulten K. VMD: Visual molecular dynamics. *Journal of Molecular Graphics*. 1996;14(1):33-38.
